# Highly-Automated, High-Throughput Replication of Yeast-based Logic Circuit Design Assessments

**DOI:** 10.1101/2022.05.31.493627

**Authors:** Robert P. Goldman, Robert Moseley, Nicholas Roehner, Bree Cummins, Justin D. Vrana, Katie J. Clowers, Daniel Bryce, Jacob Beal, Matthew DeHaven, Joshua Nowak, Trissha Higa, Vanessa Biggers, Peter Lee, Jeremy P. Hunt, Lorraine Mosqueda, Steven B. Haase, Mark Weston, George Zheng, Anastasia Deckard, Shweta Gopaulakrishnan, Joseph F. Stubbs, Niall I. Gaffney, Matthew W. Vaughn, Narendra Maheshri, Ekaterina Mikhalev, Bryan Bartley, Richard Markeloff, Tom Mitchell, Tramy Nguyen, Daniel Sumorok, Nicholas Walczak, Chris Myers, Zach Zundel, Benjamin Hatch, James Scholz, John Colonna-Romano, Lorraine Mosqueda

## Abstract

We describe an experimental campaign that replicated the performance assessment of logic gates engineered into cells of *S. cerevisiae* by Gander, *et al*. Our experimental campaign used a novel high throughput experimentation framework developed under DARPA’s Synergistic Discovery and Design (SD2) program: a remote robotic lab at Strateos executed a parameterized experimental protocol. Using this protocol and robotic execution, we generated two orders of magnitude more flow cytometry data than the original experiments. We discuss our results, which largely, but not completely, agree with the original report, and make some remarks about lessons learned.

## 1 Introduction

Replication is seen as a crisis across multiple fields of science at present, and synthetic biology is no exception. In this paper, we report results of an extensive replication experiment campaign, whose purpose was to assess the performance of novel logic gates, implemented in *Saccharomyces cerevisiae* yeast cells by Gander, *et al*. [1]. Our experimental campaign aimed to replicate the results of the original study using a very high degree of automation, and producing vastly more experimental data.

Our replication was performed using a novel, highly-automated, and high-throughput experimental framework developed through DARPA’s SD2 (Synergistic Discovery and Design) program. SD2 aims to enhance automated experimentation technology to improve replicability, experimental throughput, and experimental agility, across a range of exploratory endeavors that mix basic science and engineering, with synthetic biology as a primary area of interest. More and more laboratory automation is becoming available, increasing the scale and complexity of experiments that can be performed. Automation and information technology supports new business models with laboratory work done by technicians or outsourced to a “lab for hire.” Finally, new “multiplexing” protocols allow many tests to be conducted on a single experimental sample, and multiple experimental samples to be processed in parallel.

Prior work on replicating high-throughput experiments includes both intra- and inter-laboratory, as well as replicate-based (intra-experiment) studies. We address inter-laboratory reproducibility by comparing with the original Gander study [1] and intra-laboratory reproducibility over a number of experimental runs within the Strateos cloud laboratory. Intra-laboratory studies include work to quantify sources of variance within a protocol [2]. Important inter-laboratory studies have identified that the lack of sufficient protocol descriptions contributes to variability in plate-reader measurements [3], or proposed statistical methods to assess reproducibility [4]. Intra-experiment reproducibility via replicates is a common statistically motivated practice to better characterize samples [5, 6], and an approach used to collect our data.

Experiments in our campaign were initiated remotely, through the use of SIFT’s XPlan system [7, 8, 9], using experimental protocols captured in BBN/Raytheon’s Experimental Request framework [10] from natural language descriptions in a fixed format, stored in Google Docs. The resulting experimental requests were transmitted to a robotic laboratory operated by the Strateos company, driven by their web interface, based on the open source Autoprotocol data model [11]. After the experiments were run, measurement files were automatically uploaded to a repository hosted at the Texas Academic Computing Center (TACC), for analysis by scientists at locations throughout the US. This high-throughput workflow enables radically more experiments to be conducted, and more measurements collected.

In their paper on the design of logic gates in yeast cells Gander, *et al*. [1], describe designs for combinatory logic gates based on a core NOR gate component family (*i.e*., there are a number of different NOR gates implementable in these cells, with different gRNA used as input and output). Using CRISPR-dCas9, they build the most common six two-input logic functions out of NOR gates. Circuit diagrams and sample outputs are given in Figure 1. Their experiments explore the performance of these logic function implementations, particularly in terms of correctness of output and how clearly distinguished high and low outputs are. Providing a good “band gap” between high and low outputs is critical to enabling composition of components to compute more complex functions. Indeed, in their results circuit performance degrades with the depth of the circuit.

**Figure 1:**
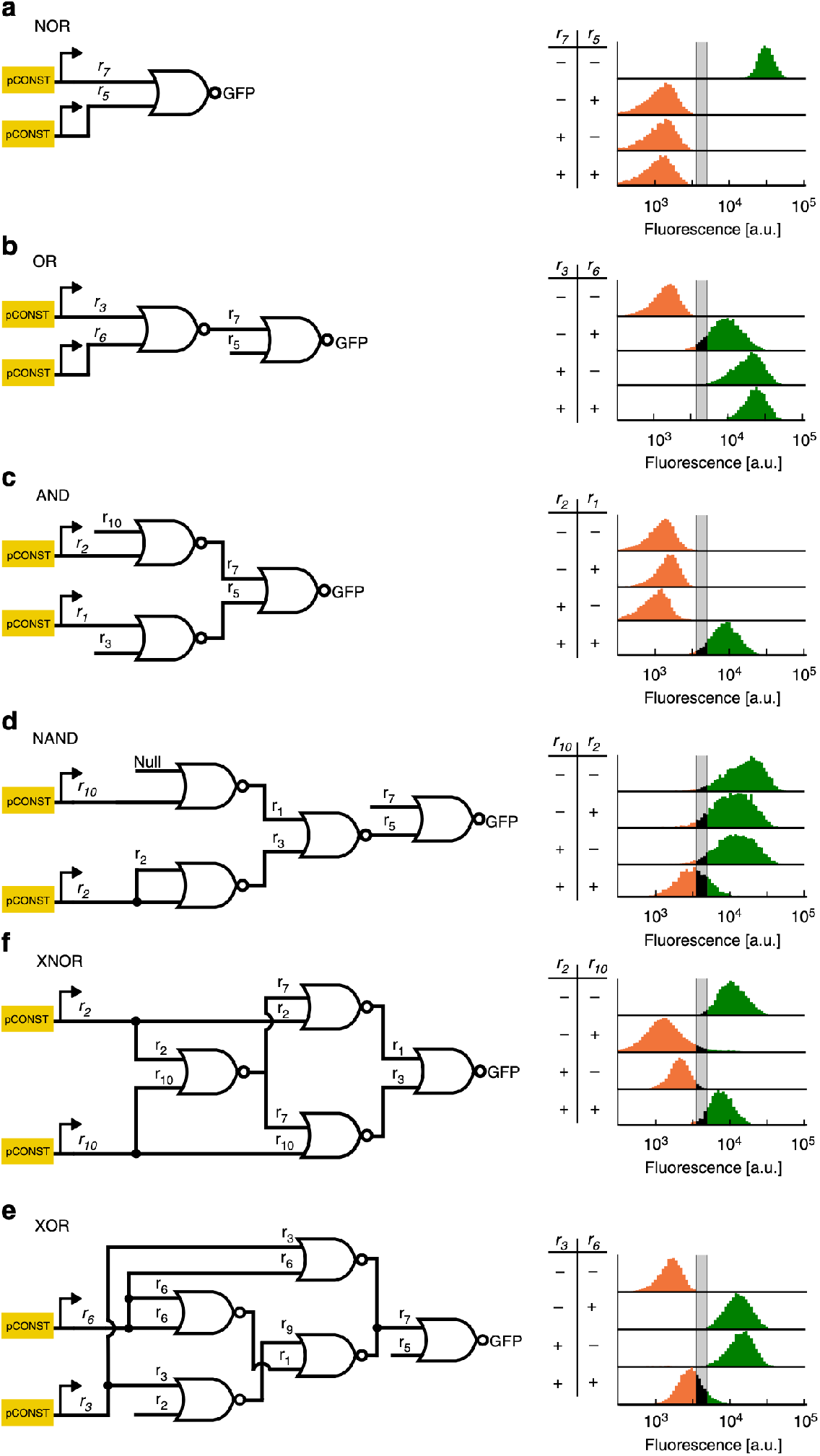
Gate structures and measured outputs, from the paper by Gander, *et al*. caption: “Six different two-input logic circuits constructed by interconnecting NOR gates. For each of the four input possibilities (−−, −+, +−, and ++), a distinct strain was constructed with the corresponding inputs expressed off of constitutive promoters (for logical +), or not integrated at all (for logical −). Fluorescence values were collected using flow cytometry of cells growing in log phase. The histograms represent population fraction from three different biological replicates measured during a single experiment and were normalized so that area sums to unity.”

The outputs of the logic functions in the yeast are measured by using the final output to produce fluorescence, which was assessed using flow cytometry (FC). Note that the gates are implemented in strains that produce their own inputs, so that there is one strain for each function × input_1_ × input_2_ combination, for example, “OR01” (or, as Gander, *et al*. sometimes write: “OR−+”).

Our experimental campaign aims to 1. replicate the original results and 2. identify factors that account for (in)correct functioning of the gates. “Correctness” of circuit function is defined in terms of the proportion of cells that exhibit high (low) output by fluorescence above Figure made available under the terms of the Creative Commons BY license. (below) a defined threshold. Growth conditions explored in these experiments include choice of growth medium, incubation temperature, target optical density (the density of the initial well populations), etc.

The experimental data we have used in the work described in this paper, with one exception, have been run in an commercial automated wet lab owned and operated by Strateos (formerly Transcriptic).Their laboratory accepts experimental requests via the internet, using a programmatic API, executes them robotically according to parameterized protocols, and then uploads the resulting data sets. The exception is the DNA sequencing data, which were collected at Ginkgo Bioworks.

Review of our results show that they generally agree with the results in the original paper. However, there are some areas of divergence. The vastly larger amounts of data available to us also show variation between individual biological replicates which are not obvious in the earlier results because of their more modest scale.

## 2 Materials and Methods

### 2.1 Experimental Strains

Strains from the original paper[1], after data collection, were stored as frozen yeast glycerol stocks at −80C. Sample ids were automatically generated and those were the ids referenced in the original paper. Strains used in the paper were derived from a strain from a collaborator on University of Washington’s campus (many years ago), which was labeled as “W303,” a commonly used laboratory strain. At the start of the SD2 program, the University of Washington Biofabrication Center (UW-BIOFAB)took the original strains from the glycerol stocks and created new cultures and glycerol stocks. From these cultures and glycerol stocks, replicate 96-well plates were created and shipped to Strateos. These were the plates used in the experiments described here.

The procedure by which these plates were frozen for storage and transport was as follows: Cell cultures were grown to log-phase in deep well 96-well plates. 20μL of each cell culture was transferred to sterilized 96-well plates (sterile 96-well PCR plates) containing 20 uL of 50% glycerol and mixed by pipetting. Plates were then covered using sterile aluminum adhesive foil and placed in insulated styrofoam containers and frozen in a −80° C freezer overnight. Plates were shipped frozen in styrofoam containers and kept frozen using dry ice.

### 2.2 Protocol

The replication experiments were executed by Strateos (previously known as Transcriptic). Strateos provides highly automated, remotely-accessible robotic wet lab services of the kind described in the introduction. Customers can specify an experiment protocol, which will be mapped onto Strateos’s lab protocol, and executed by its robotic handlers and measuring systems. In the SD2 system, these protocols are specified using Experiment Requests, which are translated and transmitted to the lab by XPLAN

The experiment campaign we discuss here involved two parameterized protocols, an initial protocol and an improved successor, the “harmonized protocol.” Parameters are indicated in the following by bold-faced capital letters (**D**, **M**, **T**, and **H**); we discuss them further at the end of this section. In this paper we analyze only the results for the harmonized protocol, but we explain the initial protocol, as well, since it produced the “standard plates” used as a starting point in runs of the harmonized protocol.

The initial protocol started with inoculating 6-well plates containing solid media from glycerol stocks of each logic gate yeast strain. A single plate contained a wild-type (WT) yeast strain, to serve as a negative control for fluorescence, and each of a logic gate’s four input states. The 6-well plates were then covered and incubated at 30C for 48 hours. After the incubation, a single colony from each well of a 6-well plate was picked and suspended into six wells on a 96-well plate containing media. A single plate contained six replicates of the WT strain and each input state of a single logic gate. Additionally, a single well on each plate was inoculated with a single colony of the NOR00 yeast strain to serve as a positive control for fluorescence. Each 96-well plate was then covered and incubated for one hour with subsequent optical density (OD) and fluorescence measurements taken via plate reader, and fluorescence measurements taken by FC. These 96-well plates were referred to as standard plates and were saved and later used in the harmonized protocol.

The initial protocol was eventually modified to better enable cross-laboratory reproducibility in the SD2 project, and the new version was named the harmonized protocol. A graphical summary of the harmonized protocol is given in Figure 2.

**Figure 2:**
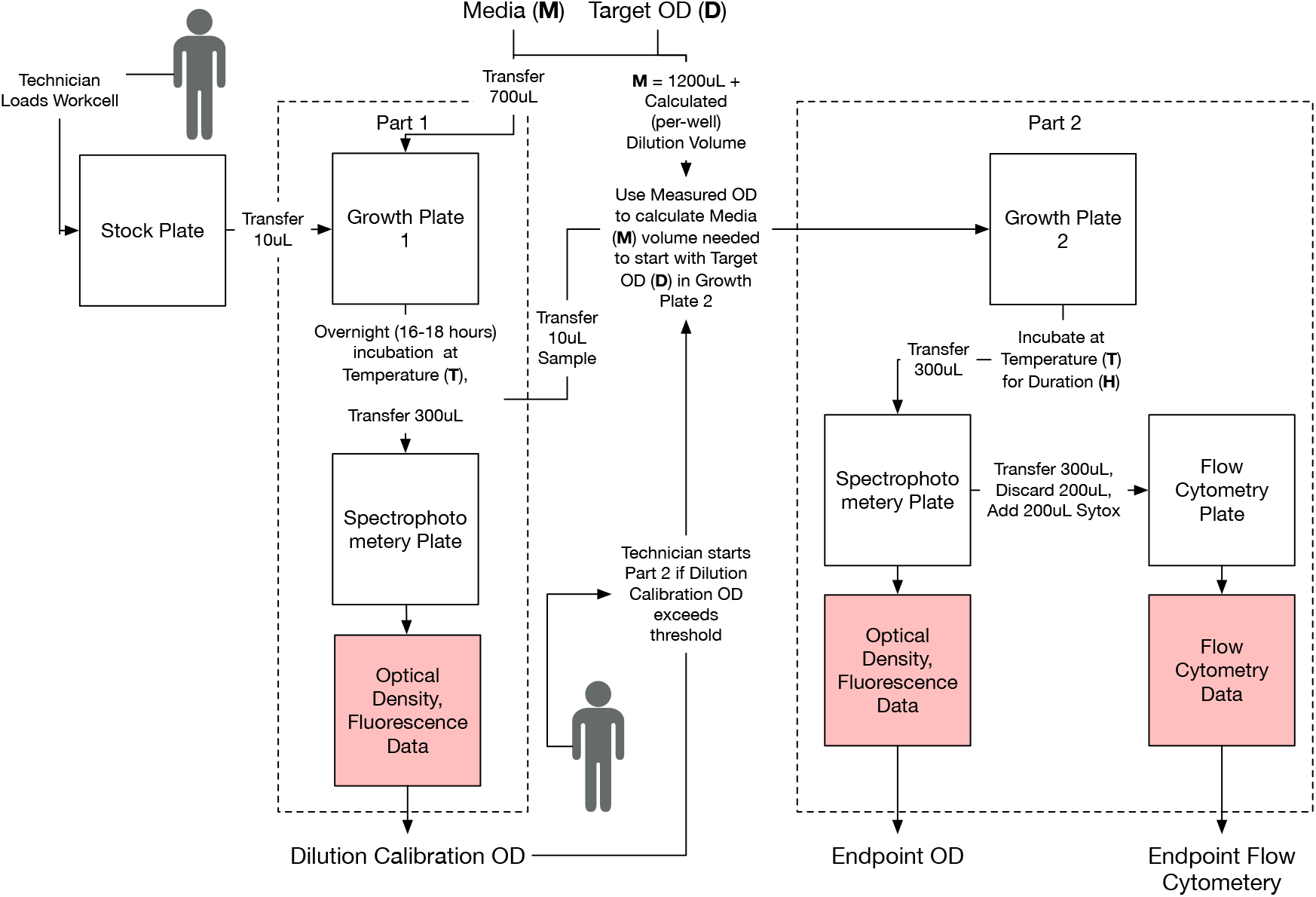
The harmonized protocol.

The harmonized protocol began with the standard plates described above, which include one aliquot of each strain, one per input state of each logic gate. Since these plates were frozen, they went through an initial “overnight recovery phase” of 18 hours (Part 1 in Figure 2). After recovery, the protocol uses a plate reader measurement to find the OD of each sample at the end of the overnight recovery phase.

The next step is to create the samples for the “growth phase” (Part 2 in Figure 2). The growth phase samples are defined by a triple (strain, target OD, replicate id). The strain is sampled from the overnight recovery phase sample plate once for each specified target OD and replicate id. The target OD, protocol parameter **D**, specifies an intended starting OD for the growth phase. For each sample triple, the protocol randomly selects a well position in the growth phase plate to hold the sample. The random assignment of well position was made out of concern that position on the plate might affect cell growth. The protocol then pipettes a sample-specific volume of growth medium (**M**) into each aliquot of the growth phase plate. The volume of medium in each aliquot is chosen so that the resulting mixture of medium and strain culture has the specified target OD. This medium volume is computed from the pre-dilution OD measured from the overnight growth plate and the target OD. For example, a sample with a measured pre-dilution OD of 1.0 can be diluted to a target OD of 0.01 by ensuring the culture to media volumes are mixed in a ratio of 1:100.

The first growth phase (Part 1 in the figure) had 700μL of growth medium and 10μL culture. The growth preparing for final measurement (Part 2) had 2000μL of media and a variable amount (approx. 10-100μL) of diluted culture (our parameter **D**). The diluted culture was 10μL culture from Part 1 and 1000μL media.

As mentioned earlier, the harmonized protocol is a parameterized protocol. One parameter was the set of strains to use in the run. The other four parameters control aspects of the growth process. The first, **T**, is the incubation temperature: this was either 30°or 37°C. We expected the higher temperature would be more challenging, since some yeast strains grow more slowly at 37°C [12, 13, 14]. The second, **H**, is the number of hours of overnight incubation, where the default was 16h, but could be varied between 8 and 18h. The third, **M**, was the growth medium used, which was either 1. synthetic complete (SC) medium (the default); 2. a rich growth medium, 3. a slow growth medium, 4. or a high osmolarity medium The high osmolarity medium was included to see if it would offset growth issues arising from the higher incubation temperature (37C). The slow growth medium was a less rich carbon source than the glucose in the standard medium. Recipes for the different growth media used are given in Supplementary Materials D. The final parameter is the target OD, **D**, which was nominally 0.0003, but varied from 1.9510^-5^to 6.3710^-1^.

Our standard growth conditions (**T** = 30°, **H** = 16, **M**= `SC` mirrored the growth conditions in the original paper to the best of our ability. They write:

Cytometry measurements were taken on cells grown in cultures diluted 1:1,000 from saturated culture for 16h at 30°C.[1, p. 9]

There is some uncertainty: we do not know what OD corresponds to “saturated” here. The growth medium is not specified here, but elsewhere (“Data collection for orthogonality matrix,” also on p. 9) specifies SC medium, and this is a reasonable assumption.

### 2.3 Laboratory Equipment

The Strateos workcell ran the harmonized protocol with the following devices: (1) Agilent Bravo liquid handler, (2) Attune NxT Acoustic Focusing Cytometer, (3) Tecan Infinite M200 Pro plate reader, and (4) Inheco ThermoShakes incubator. The protocol also used the following containers for samples: (1) Corning 96-flat (catalog #3632, 340uL wells, flow cytometry and plate reader plates), (2) Eppendorf 96-pcr (catalog #951020619, 160uL wells, stock plate), and (3) Corning 96-deep (catalog #3961, 2000uL wells, growth plate).

The harmonized protocol experiments were run in weekly batches of six to nine runs. The weekly batches were further grouped into sets of three runs that ran simultaneously, and sets were staggered over the week. Each run consisted of two parts. While run simultaneously, the runs did not necessarily represent technical replicates of the same experiment because parameterizations of the runs varied. Each run involved 93 samples (reserving 3 wells for flow cytometer calibration beads). The first part of the protocol generated one plate reader measurement per sample. The second part generated a plate reader and flow cytometry measurement for each sample. The protocol included both stamp transfers (96-to-96) and cherry pick transfers (1-to-1) between the stock plate, growth plates, dilution plates, media reservoirs, and measurement plates.

### 2.4 Protocol Analysis Methods

#### Gating

Flow cytometry measurements are designed to measure the fluorescence output from single living cells. It is a common problem that measurements of clumps of cells or cellular debris are measured, creating the need to use some type of thresholding (“gating”) to remove anomalously large or small particles from the event stream. We accomplish this through gating on forward- and side-scatter channels.

Our gating practice followed that of Gander, *et al*., who expressed a concern to avoid having budding cells in the data:

We wanted to avoid using… doublets or cells that were about to enter doublet phase because it could confound our measurement of cell state since they contain more than one copy of the nucleus and have more area to accumulate fluorescent protein. This could cause the doublets to read out a higher signal than a singlet cell. The stringent gate was designed so that we were sure that we were sampling as close to a homogeneous singlet population as we could. ^3^

We developed our gating strategy based on inspection of the positive and negative control strains, which were NOR00 and the wild type, respectively. It was not possible to simply replicate the gating used by Gander, et *al*., because the flow cytometry measurements are in arbitrary units (a.u.), and so have only relative meaning. Over the entire set of data, we found 1,831,709 FCS events associated with positive controls and 1,880,086 with negative controls. For gating purposes, we looked at both forward- (FSC_A) and side-scatter (SSC_A) FC measurements. For both positive and negative strains, heatmaps of the scatter measurements showed telltale shapes, see Figure 3.

**Figure 3:**
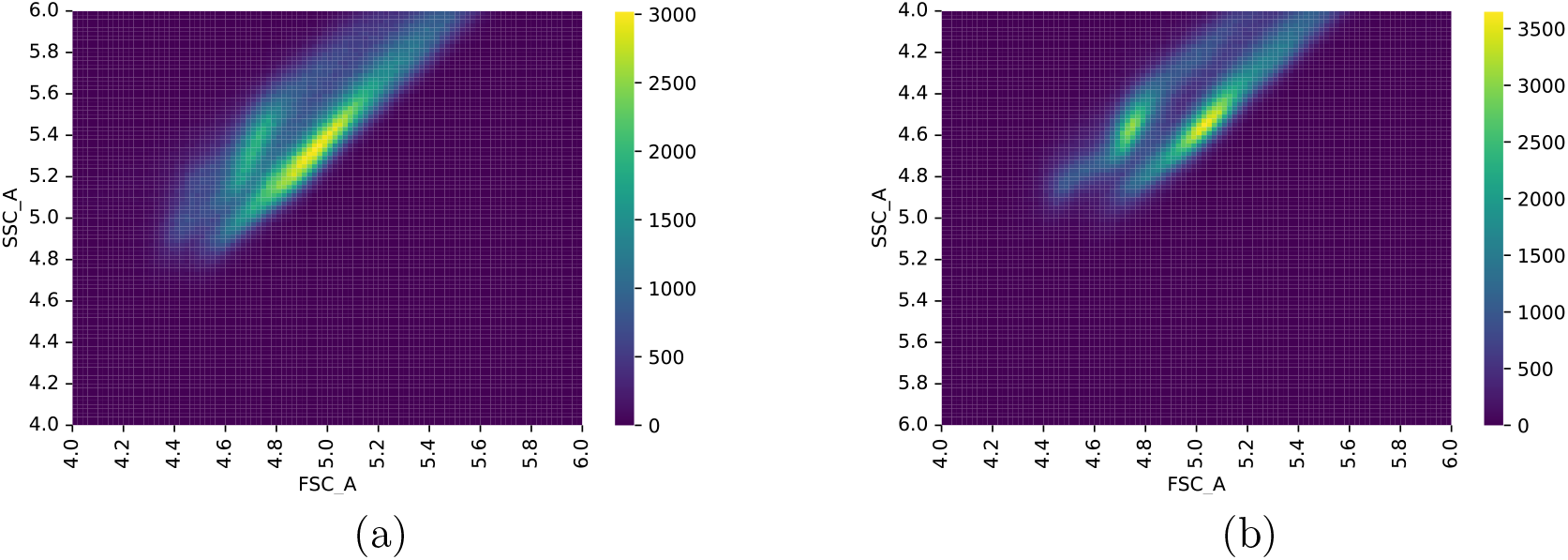
Heatmaps of scatter measurements for (a) negative controls and (b) positive controls. *x* and *y* axes are FSC_A and SSC_A measurements, respectively, plotted in log scale. Color indicates number of samples, as described in the color bars to the right of the plots.

We attempted to be conservative in the amount of possibly-useful data we discarded when gating out debris. There was also evidence that measurements were saturating at the high end, so in addition to dropping low FSC_A and SSC_A measurements, we dropped those that were exceptionally high (over 900,000 a.u., in this case). An example of this saturation is shown in Figure 4. These plots make it clear that the saturation is more pronounced in sidescatter than forward-scatter. A different illustration is given in Figure 5, these histograms vividly show the saturated measurements at the high ends, and the fact that the vast majority of the measurements (71% for the negative controls and 58.6% for the positive controls) are below 500,000 au. The flow cytometer was not sensitive to scatter measurements much above 1,000,000 au: the maximum scatter measurement for the flow cytometer was 1,048,575 au. Note that the bimodality of the responses seen in Figure 5 result from our grouping together all of the replicates for the negative and positive control strains (negative and positive controls grouped separately). We *suspect* that this is related to issues with cultures recovering from the frozen stocks: in other experiments [15] we found that on occasion recovery was slower than the 16 or 18h we allowed. We discuss this further in Sections 3.1.1 and 4.1.

**Figure 4:**
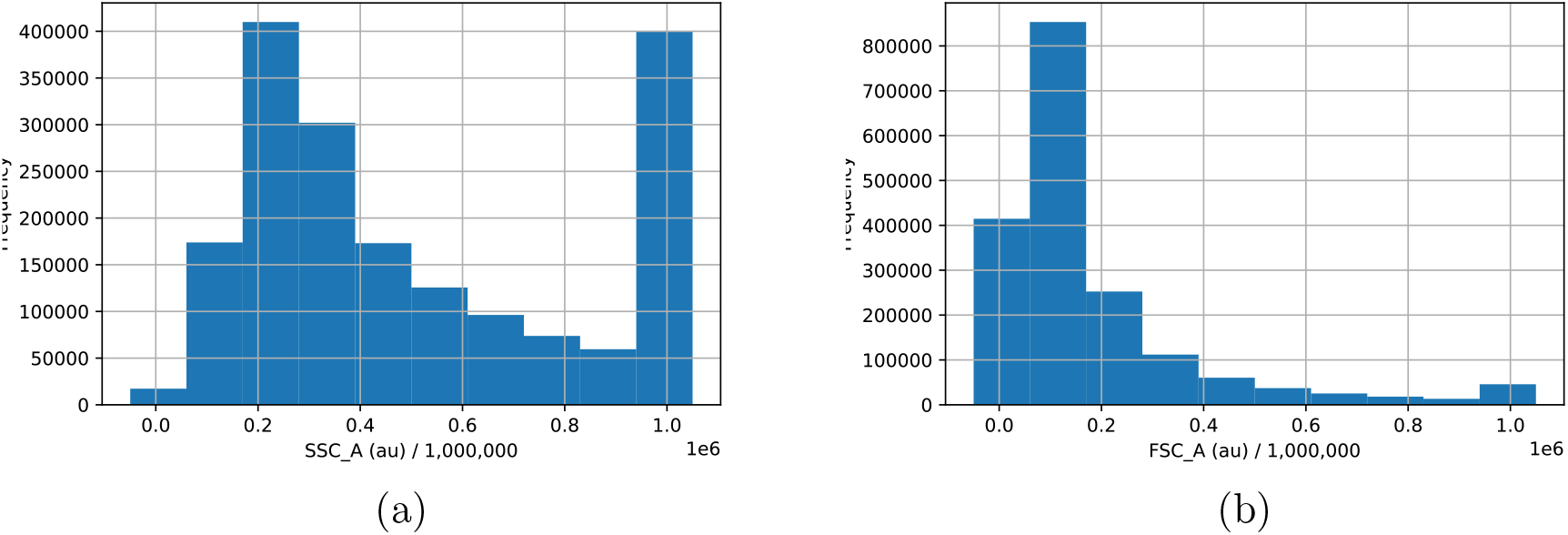
Histogram of (a) side-scatter (SSC_A) and (b) forward-scatter (FSC_A) measurements (between 0 and 1,000,000) showing saturation at the high end. Taken from positive controls.

**Figure 5:**
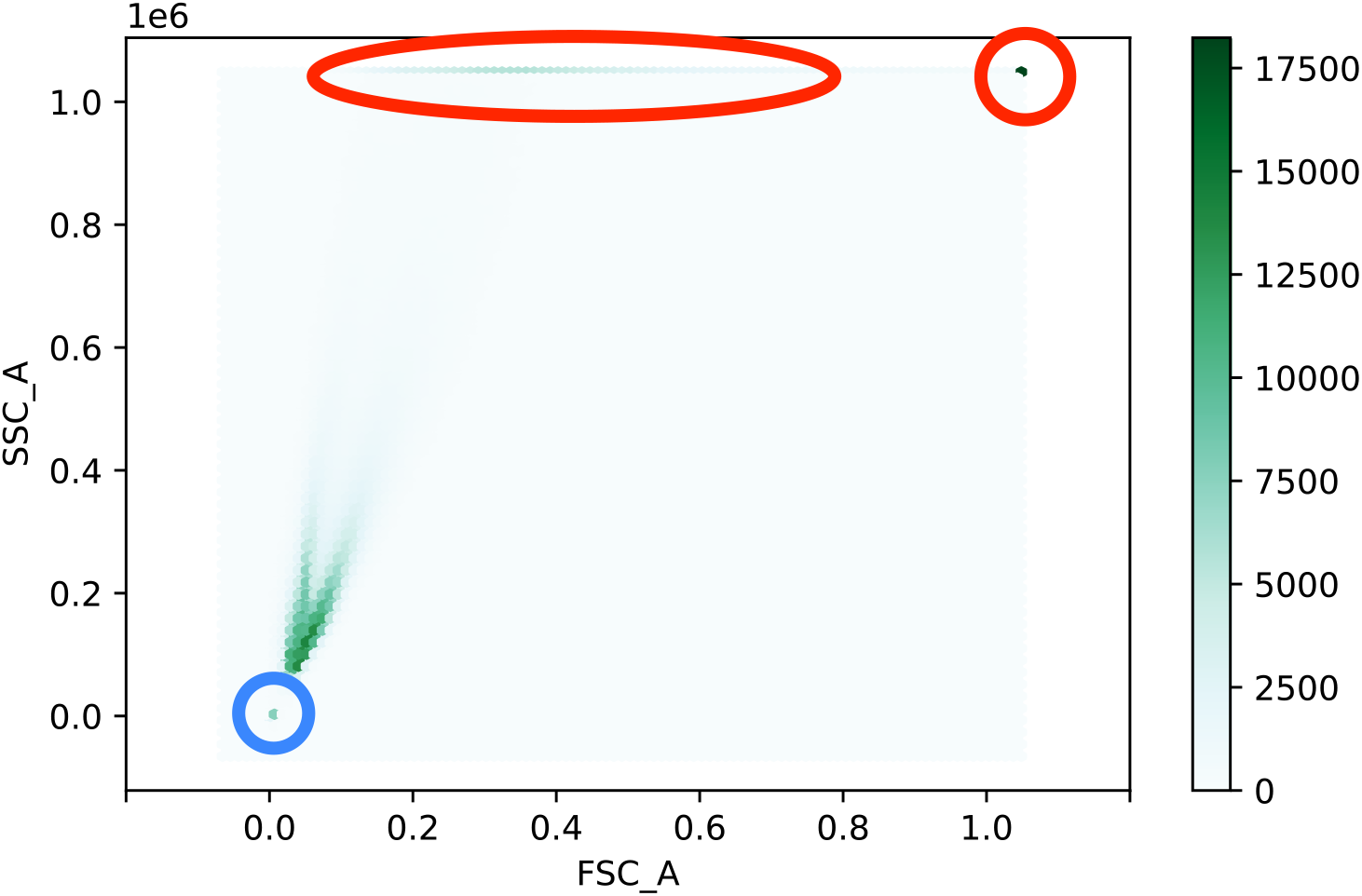
Heatmap of negative controls showing saturation effect in scatter measurements. This is an “inverse heatmap”: dark colors show regions of high concentration. Saturation at high levels is circled in red, and debris in blue. Scatter measurements are shown in fractions of 10^6^ au.

#### Eliminating Outlier Plates

We eliminated several plates that showed abnormal behavior. The process by which we identified these plates is described in Supplementary Materials E.

3 Miles Gander, personal communication.

#### FC Analysis

Flow cytometry data were aggregated at the replicate/well level, and our analyses are primarily performed based on mean and standard deviations of (gated) well GFP measurements. Our uploaded data (see Section 6) provide both raw FCS data and aggregated data frames. We decided to evaluate the FC results at the well level for both practical and principled reasons. The practical reason had to do with the difficulty of handling data from all of the raw measurement events. The more principled reason for treating the replicate/well as the unit of analysis is that (as will be discussed below), there are substantial differences between the measurements in different wells of the same strains, so that it would confound our conclusions to pool measurements across replicates.

#### Plate Reader Analysis

Plate reader data are primarily used to evaluate the density of the replicates, as a measure of their success in growing. We also use a combination of OD and fluorescence measurements as a check on the individual cell measurements from FC.

### 2.5 DNA Sequencing (DNASeq)

The DNA sequencing was conducted at Ginkgo Bioworks (also supported by the SD2 program). DNA extracted from saturated overnight cultures using a Qiagen Genomic Tip-20G kit was normalized and input to a 100x miniaturized version of the Illumina Nextera-based DNA library prep method. Final libraries were pooled, and quality control was performed via fragment analyzer (BioAnalyzer) and qPCR (Roche Lightcycler 480). The final library pool was normalized, denatured and run on the Illumina Novaseq 6000.

DNAseq analysis was performed using a Python script to search for exact matches to the gRNA DNA sequences and their reverse complement sequences in the Illumina sequencing reads for each circuit strain. The output of this script was a matrix of ones and zeros corresponding to hits and misses for the combination of each gRNA and circuit strain (the “results” matrix). A similar matrix (the “design” matrix) was constructed for the expected gRNA DNA sequences in each circuit strain by running queries against a database containing designs for each strain. A one was added to a cell in the design matrix if its row circuit design contained a target site or coding sequence for its column gRNA, and a zero was added if this was not the case. These matrices were then compared to determine whether each gRNA hit/miss was unexpected (i.e., if the value of a cell in the results matrix did not match the value of the corresponding cell in the design matrix).

## 3 Results

Before we begin, we should stress how much more data were collected in the replication campaign than in the original experiments. The original paper reports on data from three replicates (wells) for each of the 24 strains, from each of which 10,000 raw flow cytometry events were collected. After gating, there are between approximately 1000 and 3300 flow cytometry measurements per replicate (see Supplementary Materials A), for a total of 178,376 events after gating. By contrast, our highly automated pipeline made 30,000 flow cytometry measurements per replicate before gating, and over the course of the experimental campaign, approximately 8700 wells were measured. We dropped any replicate with less than 10,000 events remaining after gating (see Section 2.4) and a number of anomalous plates (see Supplementary Materials E), leaving a total of 3,923 experimental and 162 control replicates for a total of 4,085 wells of data, after gating. That yields a total of 73,928,213 experimental FC events and 3,114,226 control FC events after gating, or an overall total of 77,042,439 FC events: more than 2 orders of magnitude more than in the original experiment (see Supplementary Materials B).

### 3.1 Flow Cytometry

#### 3.1.1 Scatter results

The results of the scatter measurements that we used for gating showed anomalous results. The results we showed in our discussion of gating (Section 2.4) exhibit what looks like a bimodal distribution, a fact that we confirmed by finding two clusters in the flow cytometry scatter data using k-means clustering with *k* = 2. We see this for both positive controls (NOR11) and negative controls (Wild Type). See Figure 6. Performing linear regression on the clusters further supports the division, the *r* values of the two clusters separately being substantially better than the results of linear regression from the pooled data. See Table 1.

**Figure 6:**
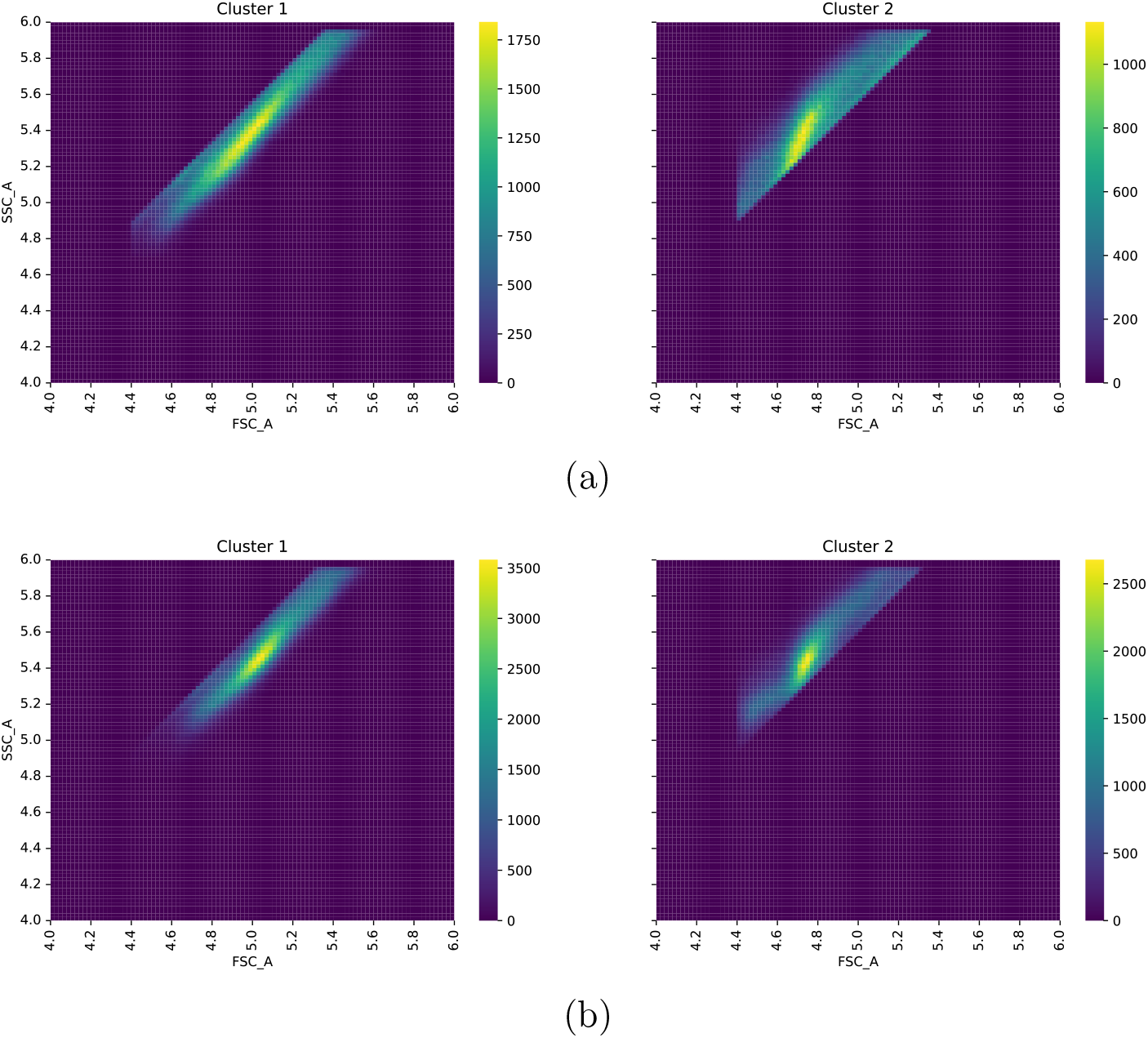
Groupings based on scatter ratios for negative (a) and positive (b) controls. Each row shows a heat map for only a single cluster. For each group, *x* axis is the log of FSC_A, and *y* the log of SSC_A. Figure was prepared by applying the k-means clustering algorithm, with *k* = 2 to the positive and negative control data. Clustering shows clear separation between the different regions shown in the heatmaps in Section 2.4.

**Table 1:**
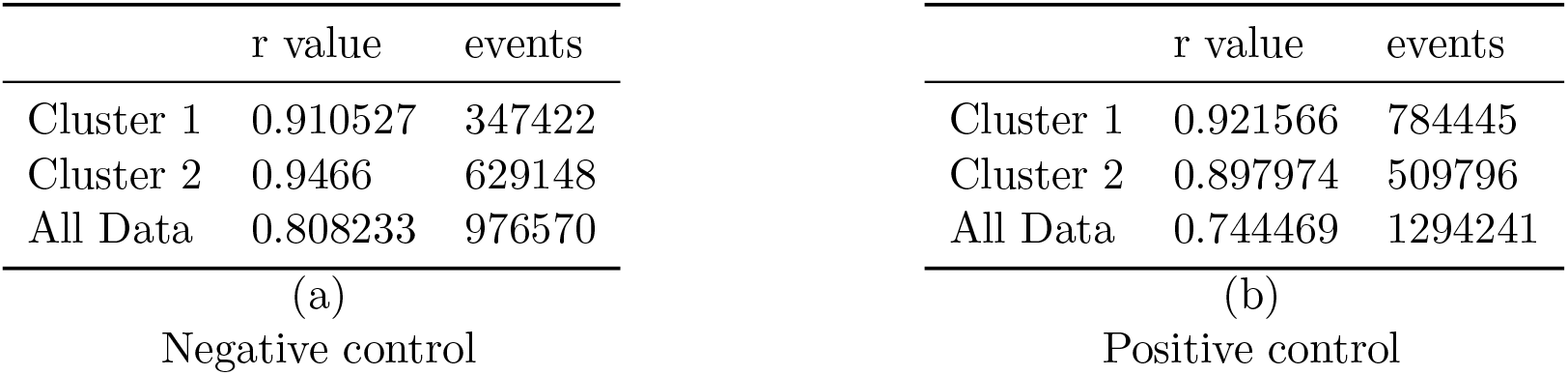
*r*-values for regression by clusters and pooled. The first two rows show the *r*-value computed for a linear regression of log(SSC_A) onto log(FSC_A) for the two clusters independently, and the third rows show the *r*-value for the same regression performed for all of the data together. We present FC event counts for the data sets, as well.

#### 3.1.2 Assessing correctness

In order to assess whether the circuits perform correctly, we must give an operational definition of correctness. Here, we propose a definition of correct functioning that takes an engineering point of view, based on a hypothetical use of these circuits in an application involving detection of some combination of environmental stimuli and computation of some response. This model suggests criteria based on performance of cell populations in aggregates: we expect biological computation to be more noisy than digital transistor circuits, so that the results of a computation will be read off as the value prevailing over a population, rather than attempting to read off an individual cell.

##### Thresholding

To operationalize this notion, we require that a replicate of a strain have a mean GFP across all events (after gating) above (below) the cutoff value to be considered correct, for outputs that should be high (resp., low). Using this definition we compute for each strain a proportion of its replicates that exhibit the correct output. Note that this definition is for replicates rather than for individual cells. For a circuit to be considered correct, all of its strains/input responses must be correct, so a circuit’s correctness is defined as the minimum proportion correct over all four of its strains/inputs. This criterion is the appropriate one, because a circuit that gives incorrect output for even one input is actually computing a *different* logic function from the one advertised. For example, under nominal conditions, the fact that the NAND gate responds incorrectly to input 11 means that the NAND gate in practice acts like a constant high output gate (see Section 3.1.3, p. 11).

The cutoff value was calculated from the set of mean gated fluorescence values for all wells, across all of the experiments, from the high and low controls. The high controls were NOR00 and the low controls, the wild type. We picked a value that optimally separated the high and low controls. The criterion for optimization was the mean distance in the “wrong direction” from the threshold: the distance above the threshold for negative control samples, and the distance below it for positive samples, divided by the number of gated control samples. The result was 236 AU, or 2.36 on the log scale.

Note that there is a value judgment made here in choosing a *single* threshold for all circuits. Again, we are driven by an engineering point of view: in order to compose together different designs, it is advantageous that they have the same threshold for interpretation as high (1) or low (0). Our choice also has the virtue of agreeing with the original paper. But this is a matter of individual preference: others might feel that it would be appropriate to choose a different threshold for each cellular logic function.

#### 3.1.3 Circuit correctness

Our analysis of the different growth conditions did not show any distinguishable effects on circuit correctness results, so we have pooled replicates from all of the growth conditions together. We give FC histograms in Figure 7. The upshot of these measurements for strain correctness is given in Table 2.

**Figure 7:**
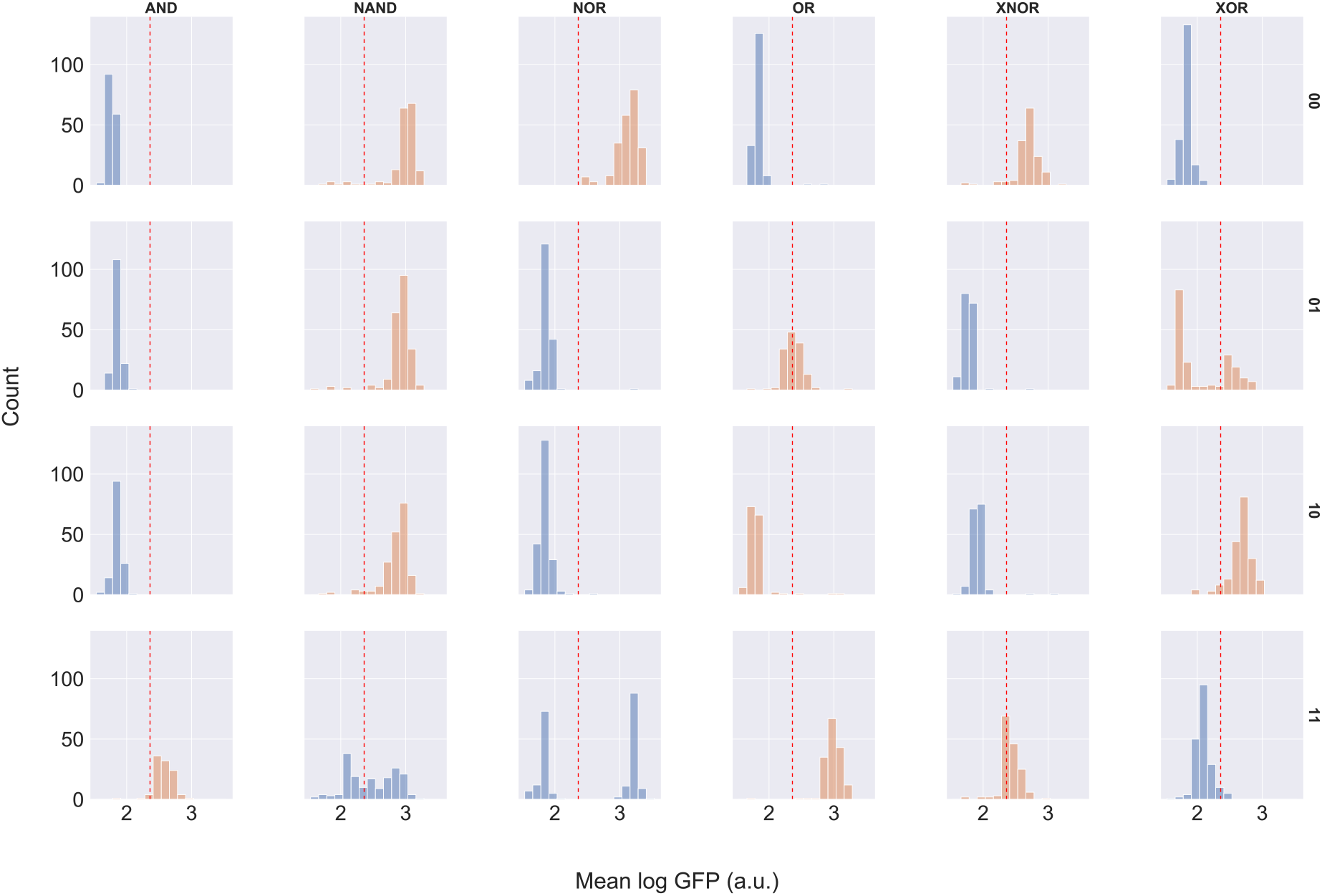
Mean GFP output per replicate for each strain, all growth conditions. Histograms are colored based on intended output, blue for strains whose outputs should be low and orange for strains whose outputs should be high. A red dashed line indicates the threshold between high and low values.

**Table 2:** Table of replicates with proportion correct. The table shows the number of replicates judged correct, “n(correct),” the total number of replicates, “n,” and the proportion correct, “p(correct).” Table cells colored gray are strains with less than 50% correct, and cells colored green are more than 90% correct.

| All Growth Conditions |  |  |  |  |
| --- | --- | --- | --- | --- |
| gate | input | p(correct) | n(correct) | n |
| AND | 00 | 0.99 | 178 | 180 |
| AND | 01 | 0.98 | 172 | 175 |
| AND | 10 | 0.99 | 166 | 167 |
| AND | 11 | 0.96 | 135 | 141 |
| NAND | 00 | 0.96 | 227 | 237 |
| NAND | 01 | 0.97 | 261 | 269 |
| NAND | 10 | 0.95 | 233 | 245 |
| NAND | 11 | 0.42 | 99 | 235 |
| NOR | 00 | 1.00 | 265 | 266 |
| NOR | 01 | 0.98 | 234 | 238 |
| NOR | 10 | 0.99 | 252 | 254 |
| NOR | 11 | 0.42 | 110 | 262 |
| OR | 00 | 0.99 | 189 | 191 |
| OR | 01 | 0.49 | 87 | 177 |
| OR | 10 | 0.02 | 4 | 188 |
| OR | 11 | 0.99 | 182 | 184 |
| XNOR | 00 | 0.95 | 200 | 211 |
| XNOR | 01 | 0.99 | 214 | 217 |
| XNOR | 10 | 0.99 | 213 | 215 |
| XNOR | 11 | 0.67 | 143 | 214 |
| XOR | 00 | 0.99 | 253 | 256 |
| XOR | 01 | 0.36 | 87 | 242 |
| XOR | 10 | 0.94 | 228 | 242 |
| XOR | 11 | 0.96 | 250 | 260 |

We have shaded the cells of Table 2 according to the proportion correct to make visual inspection easier. Cells shaded light green are those with more than 90% of replicates exhibiting correct outputs. Cells shaded gray, on the other hand, are those where less than half of the replicates exhibit correct output. The remainder of the cells are not shaded. The summary at the bottom shows that 18 of 24 strains are “green” in the full set of data, but only 11 of 24, or just less than half are “green.”

Summarizing, by our criterion that all inputs must produce the correct output, only the AND gate unambiguously works correctly, with XNOR also correct, but not as accurate in the 11 input as one would like. The NAND gate is substantially less successful: only 42% of the replicates for 11 are successfully inhibited to below the threshold value. Although the high output strains’ outputs all lie above threshold as they should, this makes the NAND gate hardly distinguishable from a constant output high. The NOR gate also performs poorly with 11 input, but handles the other inputs correctly in more than 95% of the input conditions. Interestingly, there are two distinct clusters of output values for NOR 11. For OR, neither of the single positive input conditions (01 or 10) effectively push the output high, and 10 in particular is effectively indistinguishable from 00. Finally, XOR is more than 90% correct in conditions 00, 10, and 11, but for condition 01, we see two clumps of replicates, one above the threshold, as it should be, but the majority below.

Note that the data plotted in Figure 7 are the means of **replicates**, more precisely the mean taken from gated sets of 30,000 FC events. Each data point is data for a replicate, not for a single cell. This is interesting, because apparent bimodalities, as we see in the NOR 11 strain (third column, last row of Figure 7), are not simply due to failures in individual cells: any deviation is common to the entire population in a well. This claim is confirmed by examining the standard deviation of fluorescence for each replicate. The maximum per-replicate standard deviation over all of the replicates is 0.613 logGFP (a.u.) showing that the fluorescence values of the measured yeast cells are tightly clustered, and not multimodal. Put more succinctly, lack of correctness is due to *entire wells* giving robustly incorrect values, rather than variance within wells. So it is not that wells do not provide a strong signal: instead in some cases they produce a strong *incorrect* signal, and in some other cases they produce signals that are too close to the threshold.

### 3.2 Plate reader

In addition to using flow cytometry, as in the original experiments, we also made two plate reader measurements in our protocol. These were taken 1. after a 16h recovery period of growth from frozen 96-well plates, and 2. before the strains were transferred to the experimental plates (and diluted in the process). The first measurement was taken in order to compute the dilution needed to achieve the target OD (see Section 2.2). The second measurement was taken at roughly the same time (approximately 15 minutes apart) as the FC measurements. We review these plate reader measurements now.

The two optical density (OD) readings may shed light on how successfully the various strains grow. Figure 8 shows a summary of the relationship between initial and final OD over all wells. The correlation coefficient is ≈0.44. A visual inspection reveals that there is wide variation. In addition, we observe that the final OD in a surprising number of cases is *lower* than the initial OD. There are also a substantial number of final OD values that are extremely low.

**Figure 8:**
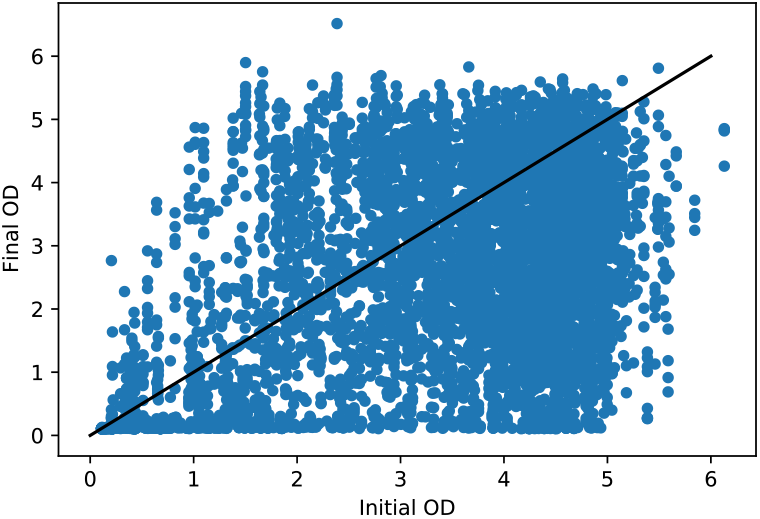
Initial versus final OD plotted for the full set of replicates.

Two obvious questions to ask are whether there are substantial differences in OD growth depending on either the growing strain, or on the incubation times. Figures 12 and 13 give this information. There is some variation by strain in the extent to which initial OD predicts final OD, and the correlation is always positive, but the relationship is weak and noisy (see Figure 12 in the Supplemental Materials F.1).

When we look at the growth relationship by incubation times, the case of 16h growth times stands out: the correlation coefficient (*ρ*) between initial and final OD for 16h recovery/16h growth is only 0.07, and for 18h recovery/16h growth, the correlation is actually *negative*: ≈ −0.15. This is in contrast to 0.28 ≤ *ρ* ≤0.46 for the other growth time conditions. All of these outlier replicates were run as part of two distinct experiment requests with identifiers `11_8_2018_1` and `2019_02_26_23_39_47`. It seems likely these are pathological in some way.

Dropping these likely problematic data sets, we find, as one would hope, a clear positive relationship between final OD and final GFP, as measured by the plate reader (see Figure 9). We conjecture that the individual bands shown correspond to the high and low outputs. Plotting the plate reader final GFP against final OD multiplied by mean log GFP from the flow cytometer shows a close agreement, as it should: the overall GFP measured by plate reader should agree with the (per cell) GFP from the cytometer multiplied by the OD from the plate reader. If these two measurements did not agree, it would signal a problem with the measurements.

**Figure 9:**
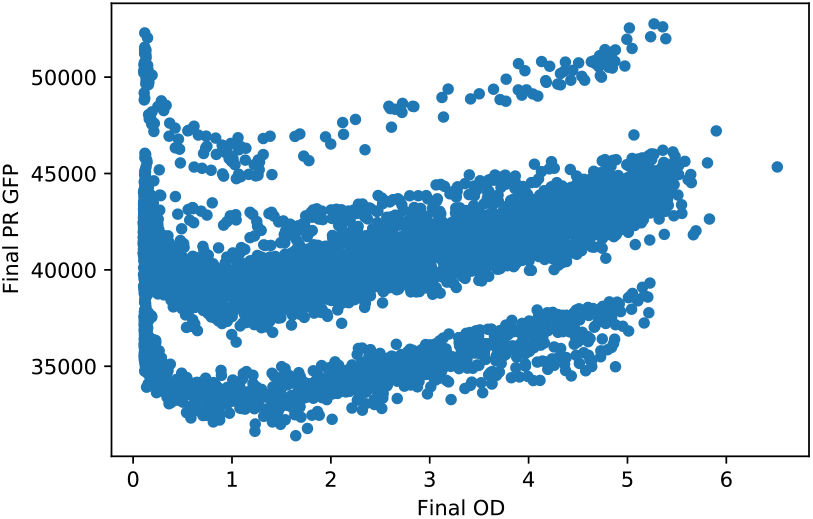
Final plate reader GFP plotted against final plate reader OD.

Finally, we regress the final OD on the initial OD (across all strains and conditions) for each setting of inoculation target OD (the **D** protocol parameter in Section 2.2, p. 5) and see that there is, as expected, an increasing relationship between the final OD and initial OD, with decreasing rate of return as **D** increases. This agrees with our background knowledge, so these results suggest that cell growth proceeded as expected. See Figure 14 in the Supplementary Materials.

### 3.3 DNA Sequencing

One of us (Roehner) analyzed DNA Sequencing data from measurements performed at Ginkgo Bioworks. See Table 3. This table shows presence and absence of gRNA DNA sequences used in the circuit designs, over a substantial subset of the experimental strains. Anomalies – sequences that were absent when the design dictated presence, and *vice versa* are highlighted. See Figure 1 to find gRNAs used in the various circuit designs. Note that this DNAseq analysis could not distinguish between whether a match to a gRNA was for a coding sequence or a target site since they have identical sequences and the analysis was performed on raw reads rather than a genome assembly.

**Table 3:** Results of searching for exact matches to gRNA DNA sequences in available raw DNAseq data for circuit strains. A cell contains 0 if its column gRNA DNA sequence is absent from its row circuit strain, and it contains 1 if its column gRNA DNA sequence is present in its row circuit strain. A cell is colored orange or blue if the absence or presence, respectively, of its column gRNA DNA sequence is unexpected based on the design of its row circuit strain. An unexpected absence of a gRNA DNA sequence may be due to an error in building a circuit strain or poor DNA sequencing coverage, while an unexpected presence of a gRNA DNA sequence may be due to a mislabeled sample, an inaccurate design specification for a circuit strain, or contamination of sequencing data for one strain with another.

| Strain | gRNA DNA Sequence |  |  |  |  |  |  |  |  |  |
| --- | --- | --- | --- | --- | --- | --- | --- | --- | --- | --- |
|  | r1 | r2 | r3 | r4 | r5 | r6 | r7 | r8 | r9 | r10 |
| AND_01 | 1 | 1 | 1 | 0 | 0 | 0 | 1 | 0 | 0 | 1 |
| AND_11 | 1 | 1 | 1 | 0 | 1 | 0 | 1 | 0 | 0 | 1 |
| NAND_00 | 0 | 1 | 0 | 0 | 1 | 0 | 1 | 0 | 0 | 1 |
| NAND_10 | 1 | 1 | 1 | 0 | 0 | 0 | 1 | 0 | 0 | 1 |
| XOR_00 | 0 | 0 | 0 | 0 | 1 | 0 | 1 | 0 | 0 | 0 |
| XOR_10 | 0 | 0 | 0 | 0 | 1 | 0 | 1 | 0 | 0 | 0 |
| XOR_01 | 0 | 0 | 1 | 0 | 1 | 0 | 1 | 0 | 0 | 0 |
| XOR_11 | 0 | 0 | 0 | 0 | 1 | 0 | 1 | 0 | 0 | 0 |
| XNOR_11 | 1 | 1 | 1 | 0 | 1 | 0 | 1 | 0 | 0 | 1 |
| NOR_00 | 1 | 1 | 1 | 0 | 1 | 0 | 1 | 0 | 0 | 1 |
| NOR_10 | 1 | 1 | 1 | 0 | 1 | 0 | 1 | 0 | 0 | 1 |
| NOR_01 | 1 | 1 | 1 | 0 | 1 | 0 | 1 | 0 | 0 | 1 |
| OR_00 | 1 | 1 | 1 | 0 | 1 | 0 | 1 | 0 | 0 | 1 |
| OR_10 | 1 | 1 | 1 | 0 | 1 | 0 | 1 | 0 | 0 | 1 |
| OR_01 | 1 | 1 | 1 | 0 | 1 | 0 | 1 | 0 | 0 | 1 |
| OR_11 | 1 | 1 | 1 | 0 | 1 | 0 | 1 | 0 | 0 | 1 |

Figure 10 shows the number of missing gRNA DNA sequence features versus an estimate of our sequencing coverage (each point represents a DNA seq data set for a strain and is colored based on the type of logic circuit sequenced). This plot was generated from the metadata automatically applied by SRA when we uploaded our data set (see Section 6). This plot does not show any evidence that low sequencing coverage was the cause of our missing gRNA DNA sequence features, but this analysis does assume that our sequencing reads are uniformly distributed across the genome and engineering constructs, which may not correspond to reality. Thus, it is possible that gRNA DNA sequence features are missing due to non-uniform sequencing coverage, but at least we can rule out individual DNA seq data sets being inadequate in terms of the total number of reads.

**Figure 10:**
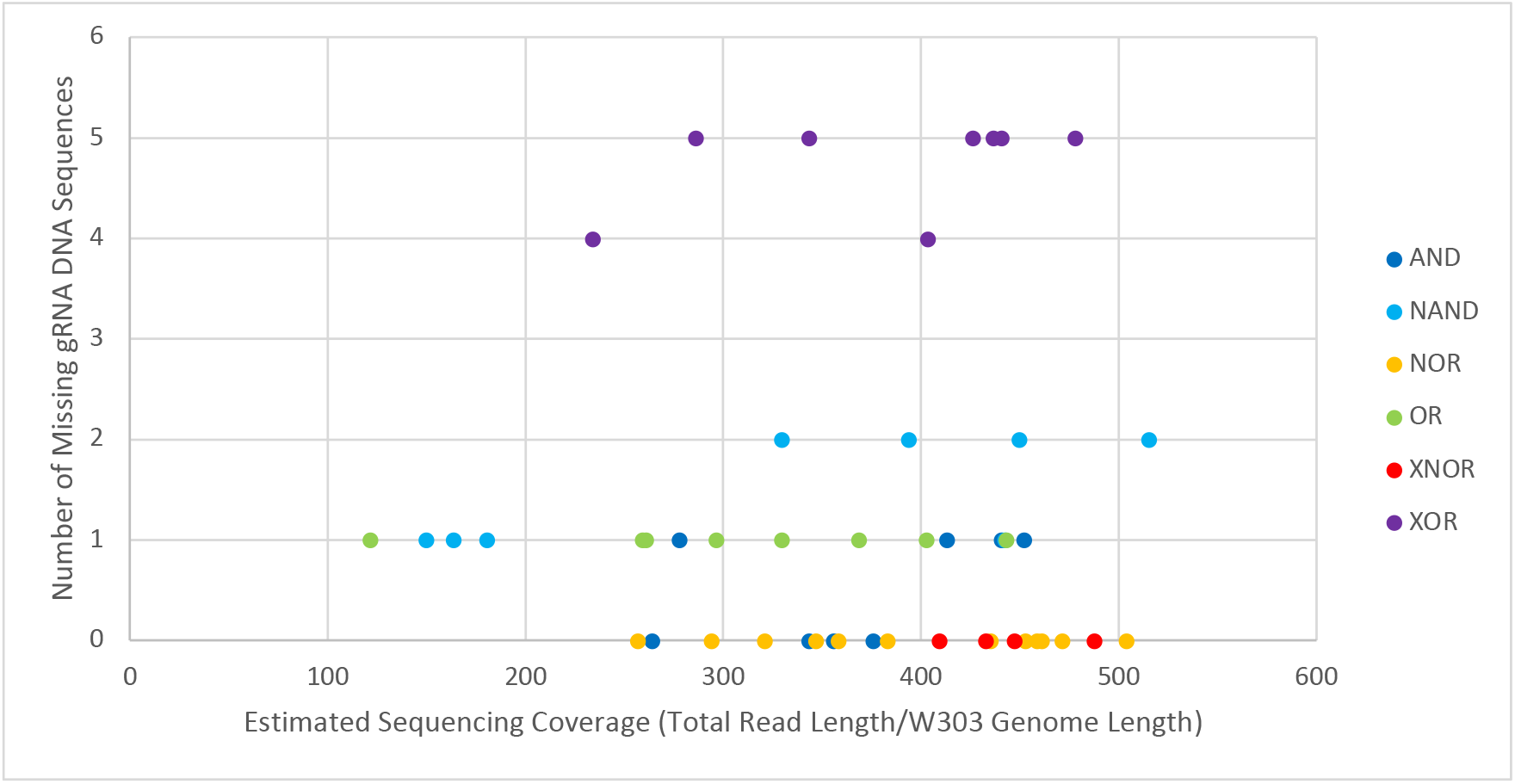
SRA Metadata analysis of our DNA Sequencing data. Sequence elements identified as missing plotted against estimated sequence coverage.

The SRA results for automated taxonomic classification of the DNA seq data (percentage breakdown of reads by kingdom, genus, species, etc.) do not show any significant differences between the data sets. They all had between 50% and 60% of their reads mapping to *S. cerevisiae* S288C, which is related to the UW BIOFAB base strain W303.

## 4 Discussion

### 4.1 Flow Cytometry Anomalies

In the process of gating the flow cytometry data described in Section 2.4, we found what looked like bimodality in the scatter measurements. In response, we further investigated the scatter data, as described in Section 3.1.1, above. Using k-means clustering, we found two distinct clusters in the plots of forward-scatter (FSC_A) against side-scatter (SSC_A). This bimodality was present in both the positive and negative control strains. We further substantiated the existence of the two clusters by comparing the results of linear regressions performed separately on the two clusters and on the pooled data (see Table 1). The correlation coefficients (r) for the linear regressions performed for the two clusters separately were substantially better than a single fit. We saw this in both positive and negative controls.

The cluster with higher ratio of side-scatter to forward-scatter could be caused by yeast cells that are either unhealthy or heading into stationary phase. This would agree with later experiments we did that showed that recovery time for these cultures was longer than we had expected [15]. We conjecture that the original cultures at BIOFAB were grown to high density, and were tending towards stationary phase when they were frozen.

It does not appear that this second cluster of values interferes with our assessment of correctness, discussed below, however. When we examine the fluorescence results for the positive controls, we do not see a substantial distinction between the two clusters. See Figure 11 for this comparison.

**Figure 11:**
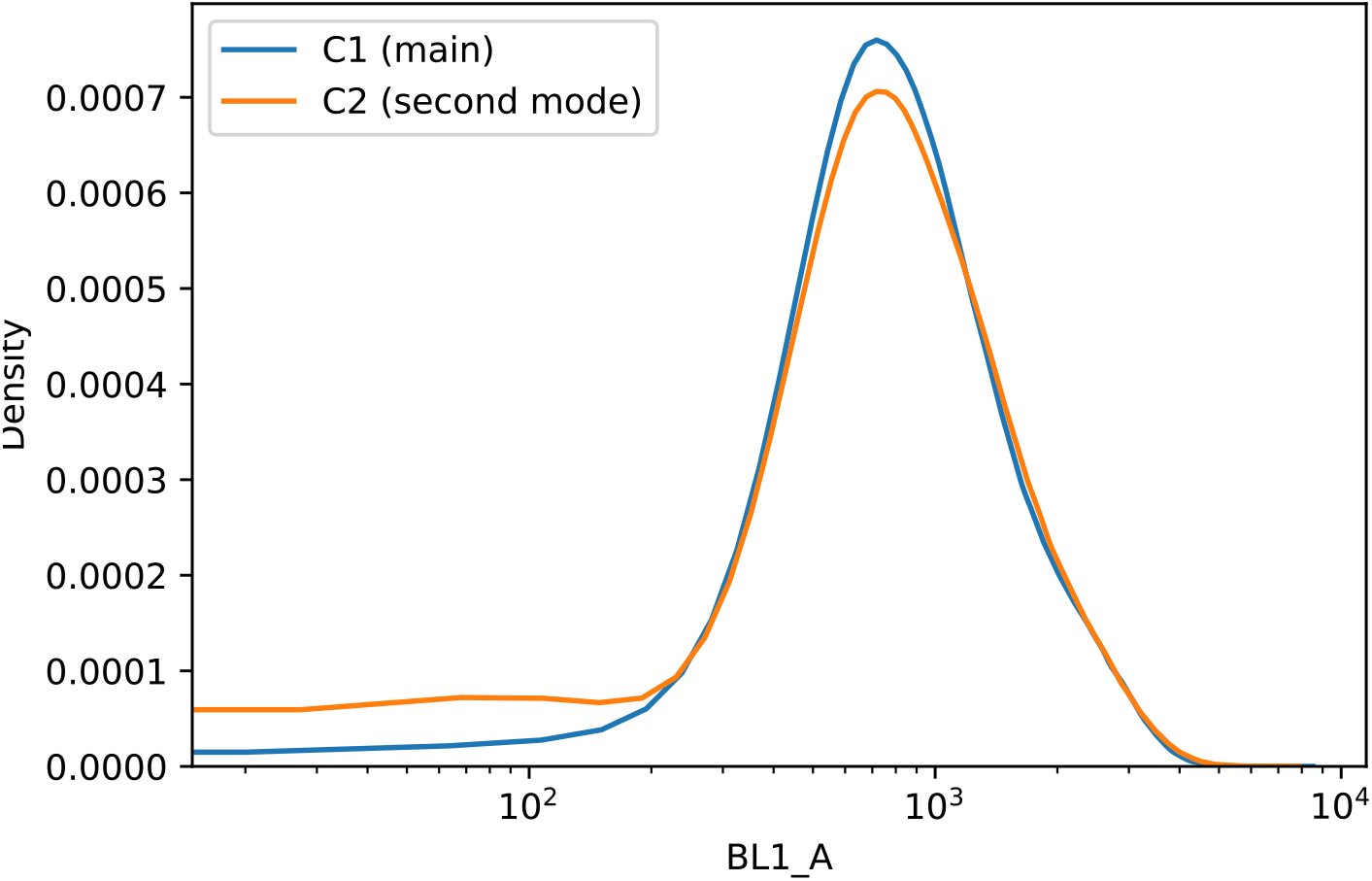
Comparing the fluorescence values of the two clusters grouped by side-scatter/forward-scatter ratio. Each curve is the KDE for one of the two clusters.

### 4.2 Plate Reader Anomalies

At first glance, it may seem anomalous that the final OD readings can be lower than the initial OD readings. However, this is only an apparent incongruity: the initial OD reading was taken in Growth Plate 1, for the purposes of choosing a dilution when transferring to Growth Plate 2. The final OD measurement comes from the descendent culture on Growth Plate 2. So while a decline in OD may show difficulty in growing, it is not an anomaly. See Figure 2 for a summary of when these measurements are taken (shaded boxes).

There is no reason to believe that variance in growth medium and culture volume led to the observed variation in growth. The pipetting equipment used is very high precision: see Section 2.3 for details.

### 4.3 Replication

In this section, we assess the extent to which our experimental results agree with those presented in the original paper. A summary of the situation, which we review below, is given in Table 4.

**Table 4:**
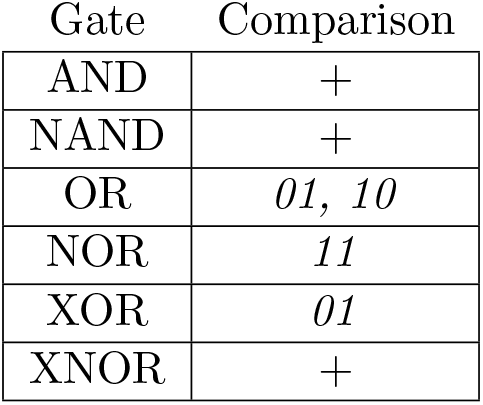
Summary qualitative comparison between original and new results. ‘+’ indicates general agreement; string indicates strain(s) with important mismatch.

For AND, our data line up well: the behavior is generally correct, but there is not as strong a separation between high and low as we would like. Similarly, comparing the original data on NAND with ours shows that for both of us, NAND failed to exhibit a clear low signal on 11: instead it straddled the threshold. There is a small difference, however: our more extensive set of results shows bimodality in the behavior of NAND11. Compare Figure 7, bottom row, first from left and Figure 1d. Note that there is a bit of an apples to oranges comparison here: the units in our graph are entire replicates, whereas the units in the original graph are individual cell measurements. XNOR looks similar: for their results as for ours, 11 is a problematic input. In the original results the signal impinges on the threshold area, but the mass of measurements is clearly above it. For us, the XNOR outputs for 11 straddle the threshold; our results look *slightly* worse, but not substantially.

For NOR, OR, and XOR, however, our data look quite different from the data in the original paper (see Figure 7 for our results). For NOR, specifically, NOR 11 shows a bimodal distribution of replicates in our experiments. It is notable that one of the conditions where there was the most difficulty in reproduction is the NOR 11 condition, since NOR is just a single isolated gate. This suggests that there may be some fragility to the dual-regulator strategy, perhaps one of the targets being less effective than the other or having an interesting constructive interference. This, in turn, would suggest this would be the place to focus on in order to improve reproducibility, as it would affect all of the other circuits too. OR is even more different – the original performance is generally quite good although the output for 01 is not as high as one might like (see Figure 1): in our results the “OR gate” behaves like an AND gate. Finally, XOR 01 is strongly bimodal in our data, although the other XOR strains behave well.

The possibility of contamination or mistakes in well positioning is a concern in light of these results. However, we do not believe that well-swapping or contamination is a likely explanation for the differences between our study and the original, partly because of the automation, and partly because we randomized the placement of strains in the wells. Contamination or well-swapping should produce a systematic effect, which we do not see in our results. Nonetheless, mislabeling during plate transfer between labs may still be an issue, as we discuss in the next section.

### 4.4 DNA Sequencing

Following up on the under-performance of the OR circuit, we noticed that the provenance of the OR strains is not clear, and that there may have been an error in the labeling or construction of one or more OR strains. As summarized in Table 3, the DNA sequencing (DNAseq) results for the OR strains did not contain the expected gRNA DNA sequences based on their design specifications. Instead, their DNAseq results more closely resembled those expected for an AND strain, which suggests that their samples may have been mislabeled. This is roughly in agreement with the flow cytometry results for OR.

This analysis revealed anomalies in several of the other strains, as well, but nothing that shows as clear a relationship to performance as for the OR gate. For example, the AND 01 strain exhibits anomalies, but the corresponding strain appears to behave well. One possibility is that the unexpected presence of gRNA DNA sequences (coding sequences/target sites) is a stronger predictor of actual problems with a circuit strain, while unexpected absences alone is more likely the result of poor DNA sequencing coverage (failure to sequence potions of the circuit strain). Three of the NOR strains show anomalies, but these are the three that appear to function correctly, and we do not have results for NOR11. XOR results are difficult to evaluate because of the complexity of the circuit.

One possibility we considered was that through some mishap, we had received a *di*ff*erent* XOR design than the one described in the original paper. In their original work, Gander, *et al*. presented 15 different designs implementing the XOR function [1, Supplement, Figure 9]. However, when we identify the gRNAs involved in these 15 implementations (see Supplementary Materials G), we see that *none* of these designs are compatible with the DNAseq results: In particular, r1 and r9 are missing from the XOR strains, but are present in every one of the designs. Similarly for r6 (which is one of the two gRNAs used as input signals).

### 4.5 Conclusions

In summary, then, our results substantially line up with the results of the original experiments. However, they also show that the performance of the current set of designs is unreliable for many gates. In follow-on work, we have extended our closed loop of experimentation to cover the design phase as well as design analysis and evaluation; we discuss these issues in two forthcoming papers [15, 16].

In the work on revising circuit designs, we were guided by results of these initial tests in deciding which circuits to redesign. The circuits redesigned were for OR and NOR. Both of these circuits showed poor performance in terms of correctness, OR being particularly bad, and in addition both had positive anomalies in DNA sequencing. Because of issues with response correctness, that follow-on work experimented with redundant designs to get more robust performance, at the expense of greater complexity and more a difficult build process. The original gates were also reimplemented from scratch, in case confusion in handling was responsible for issues in these replication experiments.

Some challenges in our work complicated the process of drawing clear conclusions. While the support of the SD2 program and the technology it developed allowed us to collect a very large amount of data to perform the replication, there was a trade off: the SD2 project was primarily a *technology development* program, and as such, the our experimental campaign was directed by the needs of the technology development, as much as anything else. Also, while the degree of automation allowed us to scale up the experiments by orders of magnitude, it came at some loss in flexibility so that, for example, we were not able to vary the temperature smoothly by degrees. We are currently working to improve the engineering of experimental protocols with an eye to making authoring and automated execution more flexible.

A final issue worth pointing out is that there could have been some confusion in the transfer between the original lab (UW’s BIOFAB) and the labs where the replication was done (Strateos and Ginkgo BioWorks). Such transfer problems are one of the many challenges to experiment replication in synthetic biology. Note, however, that all of the experiments done at Strateos were taken from the same initial plates from BIOFAB, so that any mis-labeling should be systematic *within* the work done at Strateos. What we see in the strains that perform less well looks more like stochastic failure than systematic failure.

## 5 Acknowledgements

This material is based upon work supported by the Defense Advanced Research Projects Agency (DARPA) and the Air Force Research Laboratory under Contract Nos. FA8750-17-C-O184, HR001117C0092, HR001117C0094, FA8750-17-C-0229, and FA8750-17-C-0054. Any opinions, findings and conclusions or recommendations expressed in this material are those of the authors and do not reflect the views of DARPA, the Dept. of Defense, or the U.S. Government. Approved for Public Release, Distribution Unlimited (DISTAR Case 35520, December 7th, 2021).

Thanks to Jeremy Gottlieb at SIFT for assistance with clustering the flow cytometry scatter data.

Some of the authors are employed by companies that may benefit or be perceived to benefit from this publication.

## 6 Data Availability

The DNA sequencing data are available on the NIH GEO site at this URL:https://www.ncbi.nlm.nih.gov/bioproject/?term=PRJNA784977.

The flow cytometry and plate reader data, are freely available on Zenodo with DOI 10. 5281/zenodo.6562250.

Jupyter notebooks for generating the tables and figures in this manuscript are available on GitHub at https://github.com/rpgoldman/replication-paper-data-analysis.

## Notes

https://www.ncbi.nlm.nih.gov/bioproject/?term=PRJNA784977

https://zenodo.org/record/6620957

https://github.com/rpgoldman/replication-paper-data-analysis

### Competing Interest Statement

The authors have declared no competing interest.

### Summary of Updates

Substantially improved the clarity of the manuscript. Provided more details about the protocol, notably the equipment used, and the protocols for data analysis. Also added some new plots to clarify points made in the text. Supplementary materials have been removed from the appendix and are now available in a separate file.

