## Supplementary materials for "Highly-Automated, High-Throughput Replication of Yeast-based Logic Circuit Design Assessments"

### A Sample counts from Gander, *et al.*.

All counts *after* gating.

| gate | replicate | input | count |
|---|---|---|---|
| AND | 0 | 00 | 2942 |
| | 1 | 00 | 2986 |
| | 2 | 00 | 2648 |
| | 0 | 01 | 2641 |
| | 1 | 01 | 3130 |
| | 2 | 01 | 2848 |
| | 0 | 10 | 2669 |
| | 1 | 10 | 2711 |
| | 2 | 10 | 2821 |
| | 0 | 11 | 3046 |
| | 1 | 11 | 2987 |
| | 2 | 11 | 2816 |
| NAND | 0 | 00 | 1900 |
| | 1 | 00 | 2043 |
| | 2 | 00 | 1689 |
| | 0 | 01 | 1658 |
| | 1 | 01 | 1932 |
| | 2 | 01 | 1839 |
| | 0 | 10 | 1875 |
| | 1 | 10 | 1924 |
| | 2 | 10 | 1829 |
| | 0 | 11 | 1705 |
| | 1 | 11 | 1852 |
| | 2 | 11 | 1699 |
| NOR | 0 | 00 | 2752 |
| | 1 | 00 | 2748 |
| | 2 | 00 | 2930 |
| | 0 | 01 | 2854 |
| | 1 | 01 | 2943 |
| | 2 | 01 | 3103 |
| | 0 | 10 | 3243 |
| | 1 | 10 | 3143 |
| | 2 | 10 | 2983 |
| | 0 | 11 | 2913 |
| | 1 | 11 | 3016 |
| | 2 | 11 | 3089 |

| gate | replicate | input | count |
|---|---|---|---|
| OR | 0 | 00 | 3273 |
| | 1 | 00 | 2602 |
| | 2 | 00 | 2640 |
| | 0 | 01 | 2614 |
| | 1 | 01 | 2432 |
| | 2 | 01 | 2565 |
| | 0 | 10 | 2726 |
| | 1 | 10 | 2464 |
| | 2 | 10 | 2618 |
| | 0 | 11 | 2927 |
| | 1 | 11 | 2631 |
| | 2 | 11 | 2512 |
| XNOR | 0 | 00 | 2522 |
| | 1 | 00 | 2334 |
| | 2 | 00 | 2319 |
| | 0 | 01 | 2309 |
| | 1 | 01 | 2629 |
| | 2 | 01 | 2432 |
| | 0 | 10 | 2358 |
| | 1 | 10 | 2563 |
| | 2 | 10 | 2636 |
| | 0 | 11 | 2543 |
| | 1 | 11 | 2542 |
| | 2 | 11 | 2720 |
| XOR | 0 | 00 | 2532 |
| | 1 | 00 | 2230 |
| | 2 | 00 | 2079 |
| | 0 | 01 | 2639 |
| | 1 | 01 | 2111 |
| | 2 | 01 | 2264 |
| | 0 | 10 | 2290 |
| | 1 | 10 | 2006 |
| | 2 | 10 | 2248 |
| | 0 | 11 | 2073 |
| | 1 | 11 | 1034 |
| | 2 | 11 | 1052 |

### B Sample counts from replication campaign

#### B.1 Controls

| gate | Gated count events | replicates |
|---|---|---|
| NOR-00-Control | 1,797,123 | 82 |
| WT-Live-Control | 1,317,103 | 80 |
| Totals | 3,114,226 | 162 |

#### B.2 Gates

| gate | input | Gated Counts FC events | Replicates |
|---|---|---|---|
| AND | 00 | 2,485,784 | 151 |
| | 01 | 2,535,750 | 143 |
| | 10 | 2,292,215 | 135 |
| | 11 | 2,109,267 | 101 |
| NAND | 00 | 3,133,891 | 158 |
| | 01 | 3,936,749 | 199 |
| | 10 | 3,383,250 | 172 |
| | 11 | 3,197,336 | 160 |
| NOR | 00 | 4,772,995 | 206 |
| | 01 | 3,156,527 | 175 |
| | 10 | 3,387,080 | 193 |
| | 11 | 4,106,519 | 193 |
| OR | 00 | 2,761,643 | 163 |
| | 01 | 2,643,745 | 135 |
| | 10 | 2,208,922 | 145 |
| | 11 | 3,198,050 | 155 |
| XNOR | 00 | 3,084,679 | 144 |
| | 01 | 2,442,981 | 158 |
| | 10 | 2,714,538 | 152 |
| | 11 | 3,104,314 | 149 |
| XOR | 00 | 3,111,197 | 186 |
| | 01 | 2,736,995 | 179 |
| | 10 | 3,855,776 | 185 |
| | 11 | 3,568,010 | 186 |
| Totals | | 73,928,213 | 3,923 |

### C Normal approximation plots for all (gated) replicates

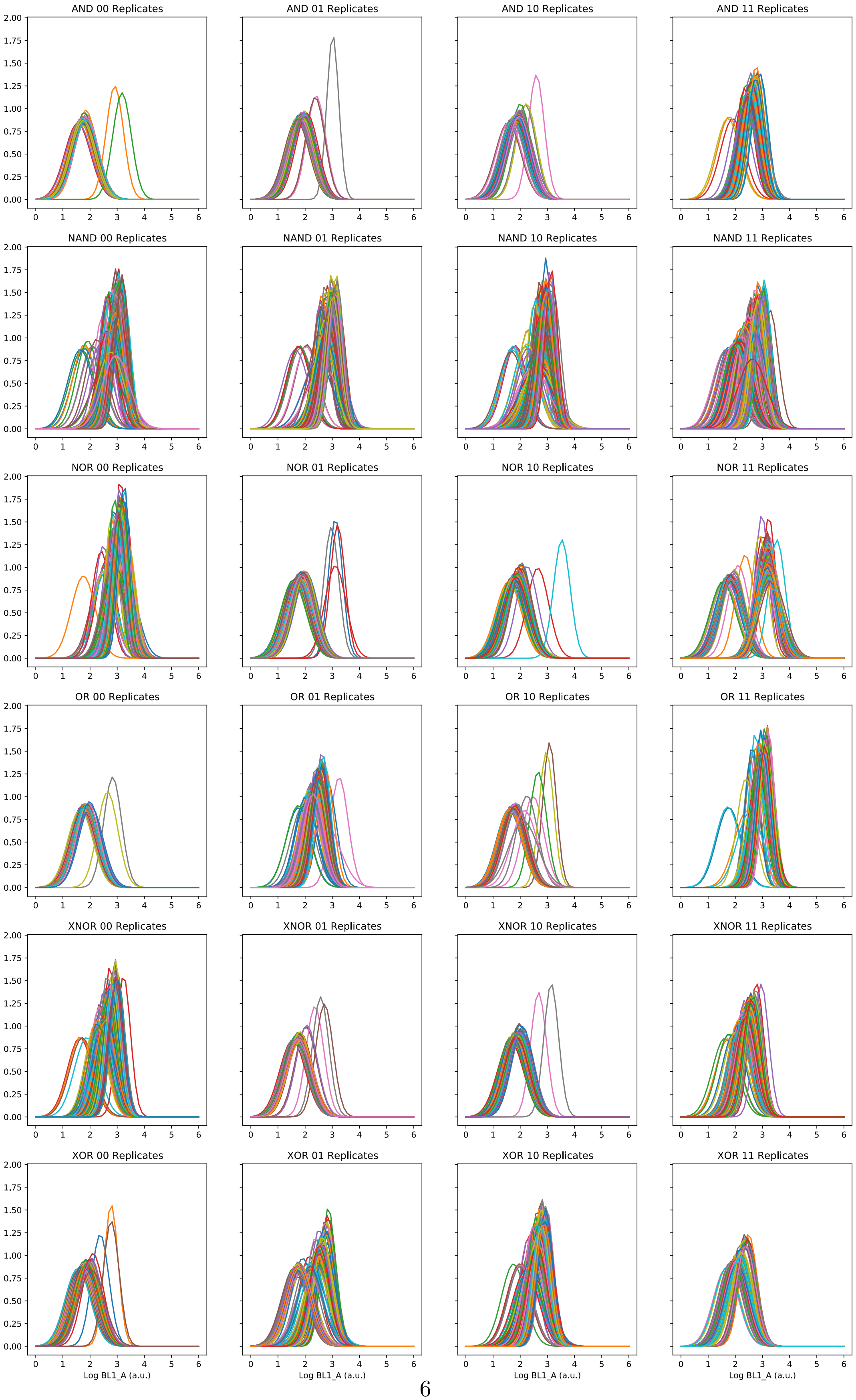

### D Growth Media

| Name | Measure | Units |
|---|---|---|
| Uracil | 0.025 | g/l |
| Adenine | 0.08 | g/l |
| ddH2O (sterile ultra-pure water) | | |
| Thermo Scientific Remel Yeast Nitrogen Base w/o Amino Acids | 6.7 | g/l |
| Dextrose (D-Glucose) | 20 | g/l |
| L-Tryptophan | 0.1 | g/l |
| L-Leucine | 0.1 | g/l |
| L-Histidine | 0.1 | g/l |
| DO Supplement -His/-Leu/-Trp/-Ura | 1.4 | g/l |

Synthetic Complete (SC)

| Name | Measure | Units |
|---|---|---|
| ddH2O (sterile ultra-pure water) | | |
| Thermo Scientific Remel Yeast Nitrogen Base w/o Amino Acids | 6.7 | g/l |
| Ethanol | 15.78 | g/l |
| DO Supplement -His/-Leu/-Trp/-Ura | 1.4 | g/l |
| Uracil | 0.025 | g/l |
| Adenine | 0.08 | g/l |
| Glycerol | 25.2 | g/l |
| L-Tryptophan | 0.1 | g/l |
| L-Leucine | 0.1 | g/l |
| L-Histidine | 0.1 | g/l |

Slow Growth (Slow)

| Name | Measure | Units |
|---|---|---|
| BD Bacto Yeast Extract BD Biosciences | 20 | g/l |
| Adenine | 0.08 | g/l |
| BD Bacto Dehydrated Culture Media Additive_Tryptone | 10 | g/l |
| Dextrose (D-Glucose) | 20 | g/l |
| ddH2O (sterile ultra-pure water) | | |

Rich

| Name | Measure | Units |
|---|---|---|
| L-Histidine | 0.1 | g/l |
| Adenine | 0.08 | g/l |
| ddH2O (sterile ultra-pure water) | | |
| Dextrose (D-Glucose) | 20 | g/l |
| L-Tryptophan | 0.1 | g/l |
| D-Sorbitol | 10 | g/l |
| Thermo Scientific Remel Yeast Nitrogen Base w/o Amino Acids | 6.7 | g/l |
| DO Supplement -His/-Leu/-Trp/-Ura | 1.4 | g/l |
| Uracil | 0.025 | g/l |
| L-Leucine | 0.1 | g/l |

High Osmolarity

| gate | input | mean | std |
|---|---|---|---|
| AND | 00 | 1.773 | 0.147 |
| | 01 | 1.854 | 0.124 |
| | 10 | 1.843 | 0.110 |
| | 11 | 2.550 | 0.165 |
| NAND | 00 | 2.945 | 0.257 |
| | 01 | 2.891 | 0.226 |
| | 10 | 2.827 | 0.237 |
| | 11 | 2.493 | 0.361 |
| NOR | 00 | 3.101 | 0.194 |
| | 01 | 1.865 | 0.182 |
| | 10 | 1.836 | 0.146 |
| | 11 | 2.606 | 0.688 |
| OR | 00 | 1.822 | 0.112 |
| | 01 | 2.362 | 0.167 |
| | 10 | 1.785 | 0.175 |
| | 11 | 2.955 | 0.185 |
| XNOR | 00 | 2.680 | 0.196 |
| | 01 | 1.755 | 0.120 |
| | 10 | 1.894 | 0.126 |
| | 11 | 2.404 | 0.167 |
| XOR | 00 | 1.824 | 0.121 |
| | 01 | 2.059 | 0.418 |
| | 10 | 2.656 | 0.179 |
| | 11 | 2.079 | 0.124 |

Table 6: Replicate means and variation thereof.

### E Removing Outliers

Close inspection of the data showed that there are several plates that have a large number of wells that are outliers. We define an outlier well as one whose gated mean GFP (by FC) value is more than 2 standard deviations from the mean value observed for that strain across all replicates in the data set (*i.e.*, the overall sample mean). See Table 6.

We found the plates that have very high numbers of outlier wells. Table 7 shows the most extreme outlier plates, those with a $p$ value of less than $10^{-6}$ (assuming normally distributed GFP values). Even with this most crude statistical analysis, it is clear that this is excess variation. Note that for two of these plates, the total well count is low, which indicates that more than 30 of the replicates were gated out (see Section 2.4). However, three of the five plates did *not* lose replicates to gating, so poor growth conditions in the wells cannot explain all of these outlier plates.

We have removed these plates from the data set used in our analyses as likely involving some form of experimental failure.

In addition to the five plates listed in Table 7, we identified another set of plates that appeared to have labeling errors: `2018_12_11_19_41_47_1`, `2018_12_11_19_41_47_2`, `2018_12_11_19_41_47_3`, `2018_12_11_19_41_47_4`, and `2018_12_11_19_41_47_5`. These have also been dropped from our analyses. These were all run very early in protocol development, which accounts for the issues.

| Plate ID | Outlier Wells | Total Wells | $p$ |
|---|---|---|---|
| r1c5vac658fxn_r1c66qw595ydy | 21 | 86 | 1.9e-10 |
| r1c7cppfr7yp6_r1c7jnv3pkbsj | 39 | 88 | 7.6e-29 |
| r1c7cprv7fe49_r1c7jmje3ebhc | 14 | 55 | 1.1e-07 |
| r1c7cpvfzqprk_r1c7fbvba55db | 27 | 87 | 8.8e-16 |
| r1c8yydkumrkr_r1c96xsxw79c9 | 20 | 52 | 4.3e-14 |

Table 7: Plates with large numbers of outlier wells.

Finally, there were two plates for which the plate reader results were suspect: `11_8_2018_1`, and `2019_02_26_23_39_47`.

### F Platereader Data

#### F.1 Initial OD versus Final OD, by Strain

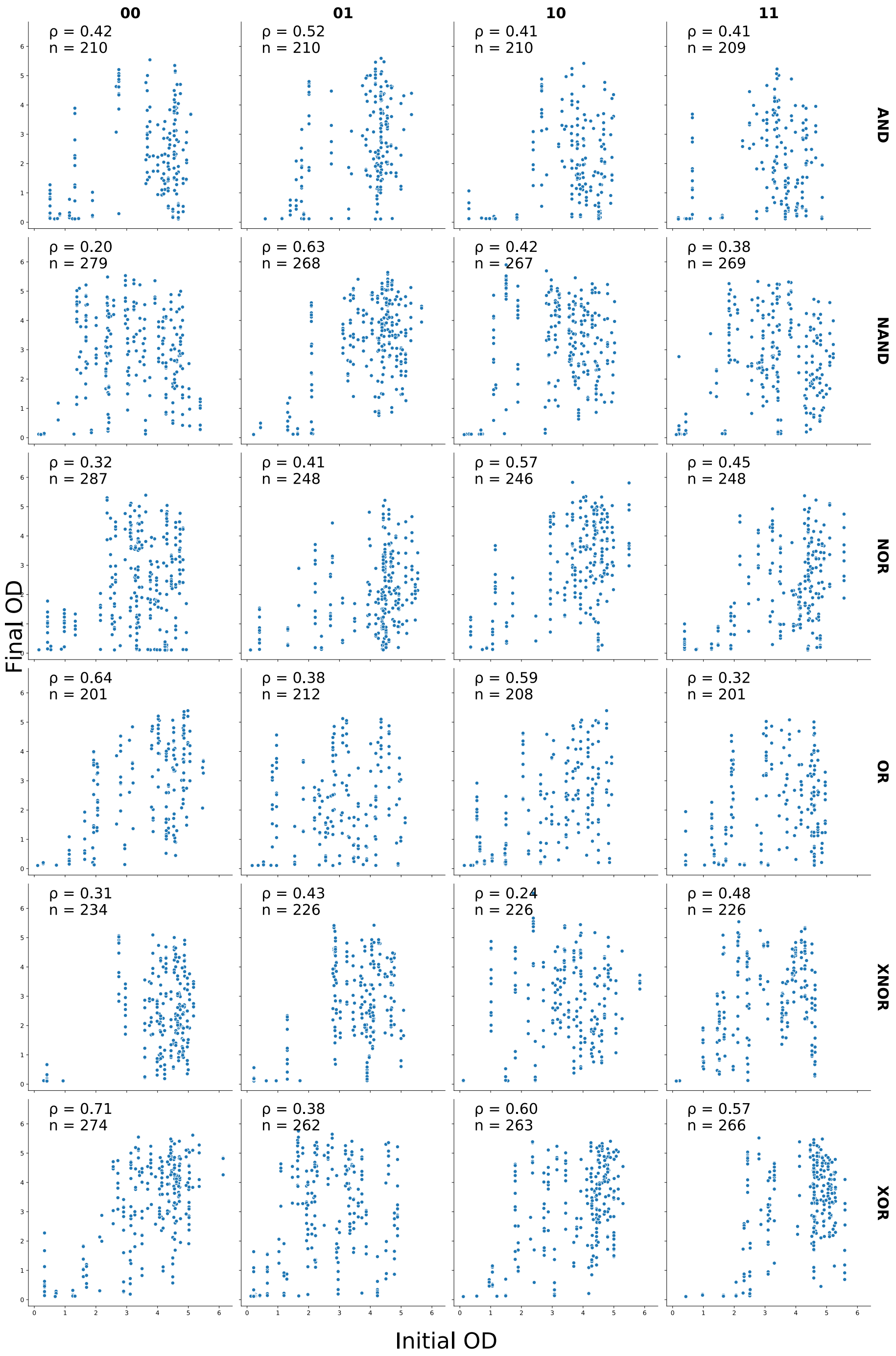


Figure 12: Growth (initial and final ODs) plotted by strain, with correlation coefficients ($\rho$) and counts ($n$).

#### F.2 Initial and Final ODs with Correlations

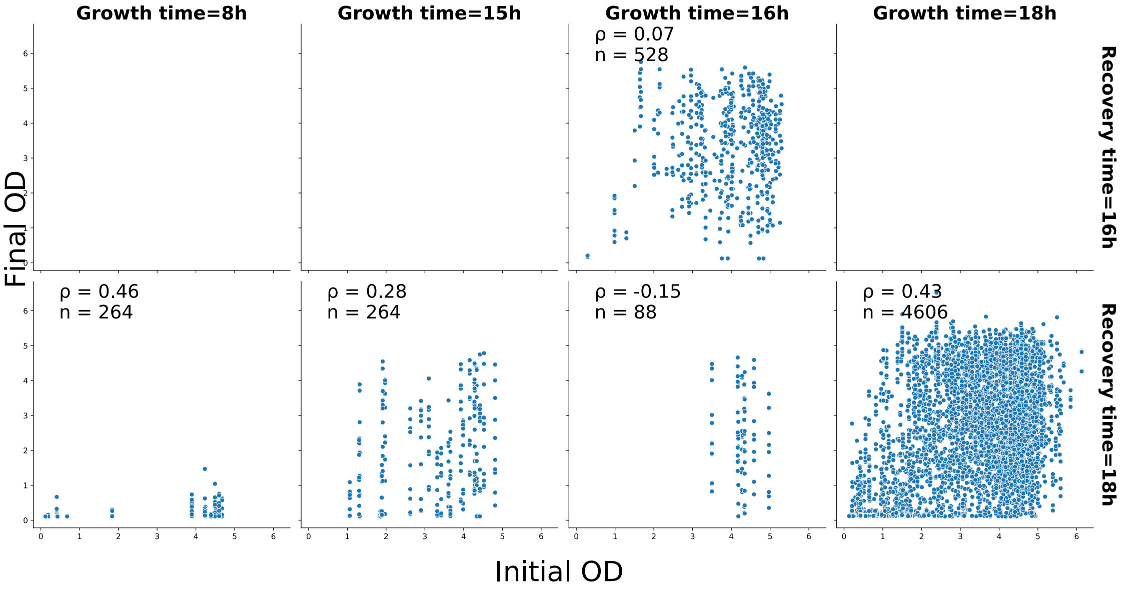


Figure 13: Initial and final ODs, plotted by growth times, with correlation coefficients ($\rho$) and counts ($n$).

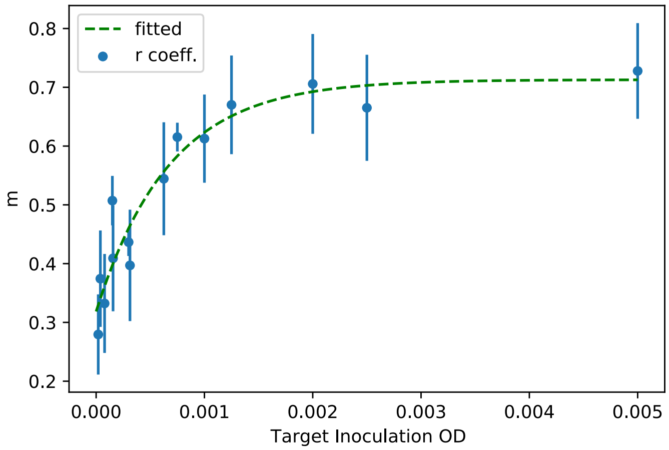


Figure 14: Regression coefficients, $m$ from final plate reader OD onto initial plate reader OD, showing increasing growth rate as a function of target inoculation OD.

#### G gRNA in Alternative XOR Designs

| gRNA present | | | | | | | | Design index |
|---|---|---|---|---|---|---|---|---|
| 1 | 2 | 3 | 5 | 6 | 7 | 9 | 10 | |
| 1 | | 1 | 1 | 1 | 1 | 1 | | 1 |
| 1 | 1 | 1 | 1 | 1 | 1 | 1 | | 2 |
| 1 | | 1 | 1 | 1 | 1 | 1 | | 3 |
| 1 | 1 | 1 | 1 | 1 | 1 | 1 | | 4 |
| 1 | 1 | 1 | 1 | 1 | 1 | 1 | 1 | 5 |
| 1 | 1 | 1 | 1 | 1 | 1 | 1 | | 6 |
| 1 | 1 | 1 | 1 | 1 | 1 | 1 | | 7 |
| 1 | 1 | 1 | 1 | 1 | 1 | 1 | | 8 |
| 1 | 1 | 1 | 1 | 1 | 1 | 1 | | 9 |
| 1 | 1 | 1 | 1 | 1 | 1 | 1 | | 10 |
| 1 | 1 | 1 | 1 | 1 | 1 | 1 | | 11 |
| 1 | 1 | 1 | 1 | 1 | 1 | 1 | | 12 |
| 1 | 1 | 1 | 1 | 1 | 1 | 1 | | 13 |
| 1 | 1 | 1 | 1 | 1 | 1 | 1 | | 14 |
| 1 | 1 | 1 | 1 | 1 | 1 | 1 | | 15 |

gRNAs present in the various alternative XOR designs by Gander, *et al.* [1, Supplement, Figure 9]. The columns shaded green are input gRNAs, and the blue are "output" gRNAs – inputs to the final NOR sub-gate that produces GFP.
